# Urbanization promotes specific bacteria in freshwater microbiomes including potential pathogens

**DOI:** 10.1101/2020.06.26.173328

**Authors:** Daniela Numberger, Luca Zoccarato, Jason Woodhouse, Lars Ganzert, Sascha Sauer, Hans-Peter Grossart, Alex Greenwood

## Abstract

Freshwater ecosystems are characterized by complex and highly dynamic microbial communities that are strongly structured by their local environment and biota. Growing city populations and the process of urbanization substantially alter freshwater environments. To determine the changes in freshwater microbial communities associated with urbanization, full-length 16S rRNA gene PacBio sequencing was performed from surface water and sediments from a wastewater treatment plant, urban and rural lakes in the Berlin-Brandenburg region, Northeast Germany. Water samples exhibited highly habitat specific bacterial communities with multiple genera showing clear urban signatures. We identified potentially harmful bacterial groups associated with environmental parameters specific to urban habitats such as *Alistipes, Escherichia/Shigella, Rickettsia* and *Streptococcus*. We demonstrate that urbanization alters natural microbial communities in lakes and, via simultaneous eutrophication, creates favorable conditions that promote specific bacterial genera including potential pathogens. Our findings are of global relevance highlighting a long-term health risk in urbanized waterbodies, at a time of accelerated global urbanization. The results demonstrate the urgency for undertaking mitigation measures such as targeted lake restoration projects and sustainable water management efforts.

## 1. Introduction

Expansion rates of urban land areas are higher than or equal to population growth rates resulting in more expansive urban land-use than compact urban growth (Seto et al., 2011). The process of urbanization leads to changes in land-cover and -use, hydrological systems, local climate and biodiversity (Grimm et al., 2008). Strong increases in urbanization are predicted over the coming decades (Gardner et al., 2016; Seto et al., 2011). While urban land areas increased by 58,000 km^2^ worldwide from 1970 to 2000, an average increase of 1,527,000 km^2^ is predicted by 2030 (Seto et al., 2011). High population density challenges freshwater hygiene and consequently human health (McLellan et al., 2015; Vörösmarty et al., 2000; Walters et al., 2011). Anthropogenic activities, such as the introduction of sewage water, and with that faecal bacteria, into natural water systems, cause eutrophication and other forms of pollution that may alter the natural microbial community composition in aquatic systems. These new microbial communities may also favour the proliferation of pathogens, whereas natural communities in nutrient-poor waters may greatly restrict such pathogen growth (Wang et al., 2013). The increasing frequency and dominance of toxic cyanobacterial blooms, but also other pathogens in parallel to anthropogenic eutrophication, pollution and warming, is of particular concern because they can directly affect human and animal health (Lapointe et al., 2015; Walters et al., 2011; Zinia and Kroeze, 2015). Further research is necessary to fully understand, how and to what extent human activities impact microbiomes of freshwater systems (Hall et al., 1999; Ibekwe et al., 2016; McLellan et al., 2015; Newton et al., 2015; Rizzo et al., 2013).

Wastewater treatment plants (WWTPs) serve the principle function of maintaining water hygiene by greatly reducing nutrients and pathogenic or harmful microorganisms (Al-Jassim et al., 2015; Asano and Levine, 1996; Numberger et al., 2019a; Wakelin et al., 2008). However, WWTPs still represent one of the major sources of freshwater pollution by dispersing pathogenic and antibiotic-resistant microbes or pharmaceuticals into the environment (Behera et al., 2011; Tahrani et al., 2015). Wastewater effluents strongly contribute to the humanization of natural microbial communities, resulting in modified water communities that contain microbes of human origin and partly resemble enteric microbiomes (McLellan et al., 2015; Newton et al., 2015; Wakelin et al., 2008). Urban lakes that are not affected by treated wastewater, remain susceptible to anthropogenic influence associated with intense recreational activity and urban storm water inflow (Newton et al., 2011; Zwart et al., 2002). Rural lakes, when not influenced by agricultural activities and other anthropogenic land-use, should exhibit natural bacterial communities, where spatio-temporal variability may be in response to environmental factors such as differences in temperature, pH, calcium carbonate and nutrient content, and organic matter availability (Allgaier and Grossart, 2006a, 2006b; Bloem et al., 1989; Güde, 1991; Newton et al., 2011; Zwart et al., 2002).

Urbanization can create conditions, which are favourable for the proliferation of pathogenic microbes. This might increase the likelihood that such pathogens may emerge (Andersson et al., 1997; Clark et al., 1996; Hunter, 2003). For example, human-introduced (micro)plastics can serve as a preferential habitat for pathogens by enabling biofilm formation in freshwater (Kirstein et al., 2016; Viršek et al., 2017). In addition, urban areas experience higher temperatures than their rural surrounding landscapes, e.g. 4.6 °C difference in the mean air temperature in the city of Beijing (Armson et al., 2012; Sundborg, 1950; Tan et al., 2010). Also, higher water temperatures are known to stimulate growth of some pathogenic bacterial species (Baker-Austin et al., 2013; Charron et al., 2004; Hunter, 2003). Moreover, in urban areas, water can be easily contaminated with pathogens by humans and pets during recreational activity (Elmir et al., 2007; Gerba, 2000; Plano et al., 2011), wildlife (Babudieri, 1958; Markwell and Shortridge, 1982), storm water runoff (Kupek et al., 2000; Schillinger and Gannon, 1985; Ward, 2002), agriculture (Givens et al., 2016; Walters et al., 2011) and wastewater effluents (Cai and Zhang, 2013; Numberger et al., 2019a; Steyer et al., 2015). Although there are hints that lake trophy and anthropogenic activity drive microbial community composition and function (Kiersztyn et al., 2019), it remains unclear which specific bacterial groups are indicative for increasing urbanization and hence may serve as indicators for environmental and human health risks.

Best practice for identifying pathogenic organisms in aquatic environments remains the utilisation of selective culture media, or molecular detection by qPCR targeting specific markers of pathogenicity (Aw and Rose, 2012; Numberger et al., 2019b). Such approaches are typically laborious, requiring multiple assays targeting distinct pathogens. Furthermore, these techniques presume a specific target and do not provide an overview on which bacteria are present. In contrast, while amplicon sequencing has been proposed as a more cost-effective method for profiling microbial communities for the presence of potentially pathogenic organisms, the obtained short-read sequences make classification to the genus level might not be that efficient than with full-length sequences (Buccheri et al., 2019). Several studies have proposed additional specific primer pairs, or increasing the number of targeted variable regions (Wang and Jia, 2016) for improved bacterial community structure determination, but these suffer from the same pitfalls for detection of bacterial pathogens. In contrast, full-length sequencing of the 16S ribosomal RNA gene, together with bioinformatic tools for determining single nucleotide variants may provide both a comprehensive profile of the microbial community and a taxonomic resolution that in many cases reaches species level (Buccheri et al., 2019; Conlan et al., 2012). Thus, this approach would offer a more precise and robust screening tool for pathogen detection in water bodies and effective monitoring of indicator species for anthropogenic pollution.

To evaluate urbanization impacts on aquatic microbial community structure, we sampled a wastewater treatment plant, three urban and two rural lakes in the Berlin-Brandenburg region (Germany) at four time points over a year. The Berlin-Brandenburg area serves as a model region as it offers a steep gradient of urbanization from a densely populated and growing city (ca. 3.7 Mio. inhabitants) to a hinterland with one of the lowest population densities in Germany (85 people per km^2^). The wastewater treatment plant treats 247,500 m^3^ raw wastewater per day generated by 1.6 million inhabitants of Berlin. The three urban lakes are classified as eutrophic and located in the German capital Berlin and the small city Feldberg in Mecklenburg-Vorpommern. The two rural lakes are located in an area in Brandenburg surrounded by forests. Their oligo-mesotrophic-eutrophic state and their undeveloped shore line together with very low population densities led us to define them as “rural lakes”. Our study combined for the first time full-length 16S rRNA gene sequencing using the PacBio Sequel I platform (Mosher et al., 2014) with the DADA2 pipeline to characterize the composition of natural microbial communities at high phylogenetic resolution, to identify bacterial groups associated with urbanization and use them as indicator species for health risk assessments.

## 2. Material and Methods

### 2.1 Sampling

The lakes Müggelsee and Weisser See in Berlin (capital of Germany with an area of 891.1 km^2^ and 3.7 Mio inhabitants) and Feldberger Haussee in the small city Feldberg in Mecklenburg-Vorpommern (pronounced anthropogenic impact due to previous wastewater input (Krienitz et al., 1996)) are comparable to lakes in bigger cities and were selected as urban lakes. None of the urban lakes received direct wastewater input during the sampling period. The two rural lakes, Dagowsee and Stechlinsee are located in a forested nature reserve area in Northern Brandenburg and have little anthropogenic impact, surrounded by only 383 inhabitants of the villages of Dagow and Neuglobsow. All lakes originate from the last ice age, but greatly vary in their present environmental status. Untreated raw inflow water and treated outflow of 1.6 million Berlin inhabitants were sampled from a municipal wastewater treatment plant (WWTP) in Berlin (Germany). This WWTP processes a negligible amount of industrial wastewater. The exact location of the sampled WWTP cannot be disclosed due to a confidentiality agreement with the WWTP operators. The characteristics of all five lakes and the wastewater treatment plant are shown in Table A.1.

Wastewater inflow and outflow, lake surface water, and sediment samples were collected every three months in 2016 from two locations in the small Lake Weißer See and three different locations in the other four selected lakes in Northeast Germany (Fig. A.1). Water was collected within a depth of 0.5 m in 2 L plastic bottles and filtered within 4 h in the lab through 0.22 μm Sterivex® filters (EMD Millipore, Darmstadt, Germany) connected to a peristaltic pump (EMD Millipore, Germany). In addition, the first centimetre of sediment was sampled using a plexiglass tube (length 50 cm, Ø 44 mm). After slicing the sediment cores in the field, samples were frozen within 2 h at −20°C until DNA extraction in the lab.

### 2.2 Measurement of nutrients and dissolved organic carbon

Due to logistic, treatment and handling challenges, measurements of environmental parameters were only successfully performed from surface lake water. Temperature and pH were measured with a digital thermometer (Carl Roth, Germany) and pH multimeter EC8 (OCS.tec GmbH & CO. KG, Germany), respectively. For measurement of orthophosphate, nitrate, nitrite, ammonium and dissolved organic carbon (DOC), 200 mL water were filtered through 0.45 μm cellulose acetate filters (Sartorius Stedim Biotech GmbH, Göttingen, Germany) after rinsing the filters with 1 mL of distilled water. The filtrate was frozen at −20°C prior to analyses. Dissolved nutrients were analysed spectrophotometrically using a flow injection analyzer (FOSS, Hilleroed, Denmark), while DOC was analysed with a Shimadzu TOC-5050 total organic carbon analyser (Duisburg, Germany). All analyses were conducted according to Wetzel and Likens (Wetzel and Likens, 1991).

### 2.3 DNA extraction

The QIAamp DNA mini kit (Qiagen, Hilden, Germany) was used for DNA extraction from Sterivex® filters (EMD Millipore, Darmstadt, Germany) following the protocol for tissue with some modifications. Prior to extraction the filters were cut into small pieces and placed into a 2 mL tube. After the addition of 200 μm low-binding zirconium glass beads (OPS Diagnostics, NJ, USA) and 360 μL of buffer ATL, the samples were vortexed for 5 min at 3,000 rpm with an Eppendorf MixMate® (Eppendorf, Hamburg, Germany). For lysis, 40 μL of Proteinase K was added and incubated at 57°C for 1 h. Then, the samples were centrifuged for 1 min at 11,000 rpm and the supernatant was transferred to a new 2 mL tube. The extraction was then continued following the manufacturer’s protocol. DNA from sediment samples was extracted using the NucleoSpin® Soil kit (Macherey Nagel, Düren, Germany), according to the manufacturer’s instructions.

### 2.4 Amplification of the full-length 16S rRNA genes

For each sample a unique symmetric set of 16 bp barcodes designed by Pacific Biosciences (CA, USA) was coupled with the primers (27F: 5’-AGRGTTYGATYMTGGCTCAG-3’ and 1492R: 5’-RGYTACCTTGTTACGACTT-3’). PCR was performed in a total volume of 25 μL, containing 12.5 μL MyFi^TM^ Mix (Bioline, London, UK), 9.3 μL water, 0.7 μL of 20 mg mL^-1^ bovine serum albumin (New England Biolabs, MA, USA), 0.75 μL of each primer (10 μM), and 1 μL of DNA. The PCR reaction included the following steps: 95°C for 3 min, 25 cycles of 95°C for 30 s, 57°C for 30 s and 72°C for 60 s with a final elongation step at 72°C for 3 min. The concentration and quality of 16S rRNA gene amplicons were measured using a TapeStation 4200 system with D5000 tapes and reagents (Agilent Technologies, CA, USA). Equimolar pools of samples were generated before sequencing. Three PCR samples containing 1 μL MilliQ water instead of sample DNA were added as negative controls and used to remove ASVs (amplicon sequence variants) representing possible laboratory contaminants from the analyzed sequences (7 unique ASVs representing 0.4% of the total sequences).

### 2.5 Library preparation, purification and sequencing

Samples were purified with the Agencourt AMPure XP kit (Beckman Coulter, USA) and sequencing libraries including DNA damage repair, end-repair and ligation of hairpin adapters were prepared using the SMRTbell Template Prep Kit 1.0-SPv3 following the instructions in the amplicon template protocol (Pacific Biosciences, USA). The Sequel Binding Kit 2.0 (Pacific Biosciences, USA) was used to bind DNA template libraries to the Sequel polymerase 2.0. The data for each sample were collected in a single Sequel SMRT Cell 1M v2 with 600 min movie time on the Sequel system I (Pacific Biosciences, USA). The Diffusion Loading mode was used in combination with a 5 pM on-plate loading concentration on the Sequel Sequencing Plate 2.0 (Pacific Biosciences, USA). The SMRT Analysis Software (Pacific Biosciences, USA) generated Circular Consensus Sequences (CCS) for each multiplexed sample that was used for further downstream analyses.

### 2.6 Bioinformatics and statistics

All 13 SMRT cells together generated 2.4 M subreads with an average length of ~25 kb and a mean of 17 passes of the 16S rRNA gene (expected length of ~1.5 kb). Circular consensus sequences (CCS) for multiplexed samples were generated from the subreads using pbcss v3.4 (https://github.com/PacificBiosciences/ccs) with -minPredictedAccuracy of 0.99 and sample demultiplexing was performed using lima v2.0.0 (https://github.com/pacificbiosciences/barcoding/) with the parameters --same, --ccs and --min-score 80. Removal of primers was carried out using Cutadapt (Martin, 2011) while additional quality filtering, trimming, and identification of unique amplicon sequence variants (ASVs) was performed using the DADA2 pipeline in R (Callahan et al., 2016, p. 2). As DADA2 relies on a minimum amount of sequence replication to infer the “true” sequence variants, we had to further trim the CCS because when pooling all ~2.4 M CCS together, only 1.6% of the original CCS was found to be duplicated. CCS were trimmed using different combinations of hypervariable primers of the 16S rRNA gene aiming to retain the highest number of sequences in the final ASV table while losing the minimum number of bases (Table A.2). The CCS were trimmed at the 5’ end using the forward V2 primer (ACTCCTACGGGGAGGCAGCA) which led to an average length of 1,123 bp (versus 1,443 bp of the untrimmed CCS), 4.4% of sequence duplication and the retention of 2 M sequences in the final ASV table (Table B.1). The taxonomic classification of the ASV table was performed with SINA v1.7.2 against the SILVA reference database (SSU NR 99 v138.1) (Pruesse et al., 2012; Quast et al., 2013; Yarza et al., 2014) and ASVs assigned as “Chloroplast” removed. All down-stream analyses were performed in R v4.1.0 (R Core Team, 2013).

Weighted correlation network analysis (WGCNA package (Langfelder and Horvath, 2008)) was carried out to identify modules of the bacterial community enriched in potential pathogenic taxa. Briefly, any noisy signal from rare ASVs was removed from the ASV table by retaining only ASVs which occurred with 10 or more sequences in at least 3 samples (this step removed 4670 and 3287 ASVs, representing 9.5% and 20% of total sequence abundance in water and sediment samples, respectively). An adjacency matrix was computed using the function *adjacency* on the centred log-ratio transformed ASV sequence counts (*clr* function, package compositions; (Parent and Parent, 2015)) to ensure sub-compositional coherence. The function infers ASVs connectivity by calculating an ASV similarity matrix (based on Pearson correlation) and applying soft thresh-holding to empathize the strongest correlations. The soft threshold value 5 was picked with the function *pickSoftThreshold* as it was the smallest value achieving a R^2^ > 0.9 for a scale-free topology fit. Topological overlap dissimilarity was calculated with the function *TOMdist* on the adjacency matrix and fed into a hierarchical clustering (*hclust* function, Ward.D2 agglomeration method). ASV modules were automatically identified on the clustering by mean of the function *cutreeDynamic* to identify branch boundaries for modules (deepSplit = 4 and minClusterSize = 20). The ASV modules were summarized by their first principal component (function *moduleEigengenes*) which was correlated with vectors of relative abundance of the potential pathogenic groups to identify ASV modules significantly enriched in those groups. The vectors of potential pathogens were obtained by summing the relative abundance of the ASVs classified as belonging to either one of the potential pathogenic taxa (according to NCBI Pathogen Detection Project, see http://www.ncbi.nlm.nih.gov/projects/pathogens/) across all samples and transformed using centred log-ratio. Enrichment and *p*-values were obtained from a univariate regression model between each module principal component and each vector of potential pathogens and results were visualized as a heatmap.

Ternary plots were plotted/generated using the function *ggtern* (package ggtern; (Hamilton and Ferry, 2018)) while an indicator species analysis was carried out by mean of the *multipatt* function (func = *“IndVal.g”,* duleg = F, max.order = 2; package indicspecies) to identify ASVs specifically associated to each of the habitats (i.e. wastewater, urban and rural lakes) and their combination (Cáceres et al., 2010; Cáceres and Legendre, 2009). Only ASVs with a p-value adjusted for false discovery rate < 0.05 were considered.

Non-metric multidimensional scaling (NMDS) analyses were generated by the function *metaMDS()* using package vegan in R version 3.5 and Bray-Curtis dissimilarity index. Constrained correspondence analyses (CCA) were also performed in R using the package vegan and the function *cca()* followed by an one-way analysis of variance (ANOVA) with the function *anova()* and ‘n perm=999’ (Dixon, 2003; Oksanen et al., 2014; R Core Team, 2013). Boxplots were created by using the R packages ggplot2 and ggsignif with Wilcoxon test for the significance levels (Ahlmann-Eltze, 2017; Wickham, 2009).

## 3. Results

### 3.1 Between and among lake bacterial community heterogeneity

Surface water samples were dominated by Gammaproteobacteria (32.9 ± 13.2%), Cyanobacteria (14.9 ± 18.7%), Bacteroidota (11.7 ± 5.8%), Actinobacteriota (11.4 ± 8.2%) and Alphaproteobacteria (10.8 ± 4.9%), Planctomycetota (8.8 ± 8.0%) and Verrucomicrobiota (5.7 ± 3.1%). The most abundant phyla in the sediment samples were Gammaproteobacteria (44.1 ± 7.9%), Bacteroidota (16.7 ± 4.5%), Alphaproteobacteria (8.2 ± 3.4%), Cyanobacteria (5.1 ± 4.2%), Verrucomicrobiota (6.0 ± 2.9%), Planctomycetota (3.4 ± 1.8%) and Acidobacteriota (3.1 ± 1.1%).

Non-metric multidimensional scaling (NMDS) analyses of the lakes were performed to display seasonal or spatial heterogeneity within a lake. In each lake two main clusters could be defined: water and sediment. Furthermore, water samples showed a higher variance than the sediment samples, which were more similar to each other, except for Müggelsee. Among the water samples we observed a clustering of samples by season, whereas the sediment samples revealed either random (no clear pattern) or spatial patterns according to the sampling site (Fig. A.2).

Environmental parameters and nutrients were measured in surface water samples and used to detect any significant correlations with the bacterial communities/groups. A constrained correspondence analysis (CCA) of the surface water samples in combination with an analysis of variance (ANOVA) showed that temperature, pH, orthophosphate, nitrate, nitrite, ammonium and dissolved organic carbon (DOC) concentration had a significant (all *p*≤0.001) correlation with the composition of the lake bacterial communities (**Fig. 1a**). Overall, temperature (χ^2^ = 0.4219) and orthophosphate concentration (χ^2^ = 0.3594) had the strongest correlation with the community composition. Nitrate, orthophosphate and temperature displayed a stronger correlation with urban lake than with rural lake samples. Among the most abundant bacterial phyla, Alphaproteobacteria and Bacteroidota showed a positive correlation with temperature. However, the CCA only explained 10.7% of the total variance (**Fig. 1b**).

**Figure 1:**
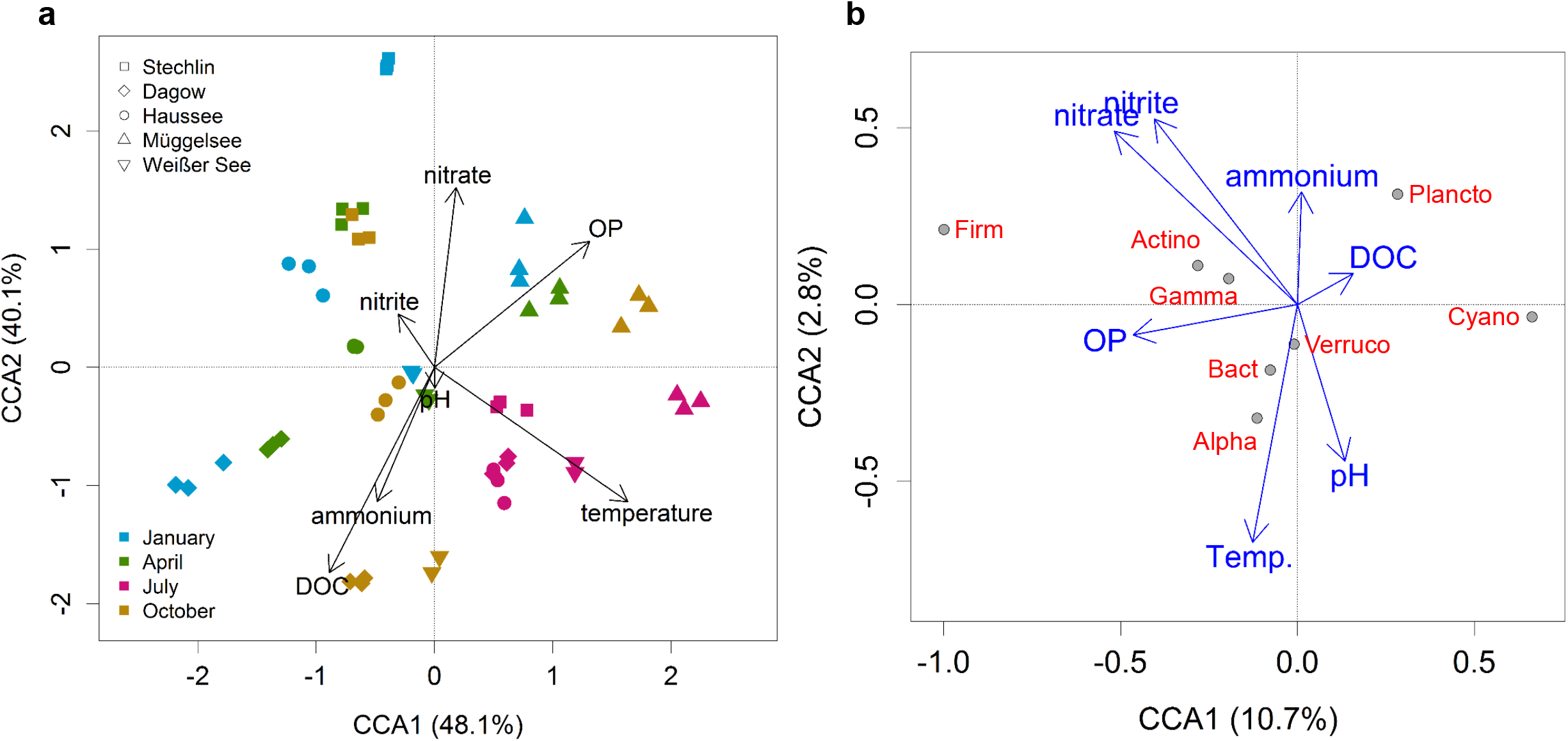
Influence of environmental parameters on the bacterial communities in surface water. **[a]** A constrained correspondence analysis (CCA) of all water samples and their corresponding environmental measurements: ammonium, dissolved organic carbon (DOC), nitrite, nitrate, orthophosphate (OP), pH, and temperature is shown. Colours indicate the season (blue: winter, green: spring, pink: summer and brown: autumn) and the symbols show the different lakes (square: Stechlinsee, diamond: Dagowsee, circle: Feldberger Haussee, triangle: Müggelsee and inverse triangle: Weißer See). **[b]** A constrained correspondence analysis (CCA) shows the most abundant bacterial phyla Actinobacteriota (Actino), Alphaproteobacteria (Alpha), Bacteroidota (Bact), Cyanobacteria (Cyano), Firmicutes (Firm), Gammaproteobacteria (Gamma), Planctomycetota (Plancto) and Verrucomicrobiota (Verruco) and their correlations with the environmental measurements in lake surface water samples.

### 3.2 Habitat-specific bacterial communities in rural and urban freshwater habitats

Lakes were generally characterized by significantly higher fractions (relative abundance) of Actinobacteriota, Alphaproteobacteria, Planctomycetota and Verrucomicrobiota, while wastewater had significantly higher levels of Firmicutes and Gammaproteobacteria (without the order Burkholderiales) (**Fig. 2**). Urban lakes differed significantly from rural lakes by having higher relative abundances of Actinobacteria, Alphaproteobacteria, Burkholderiales and Firmicutes. The Bacteroidota were significantly higher in the wastewater outflow and lakes than in the wastewater inflow.

**Figure 2:**
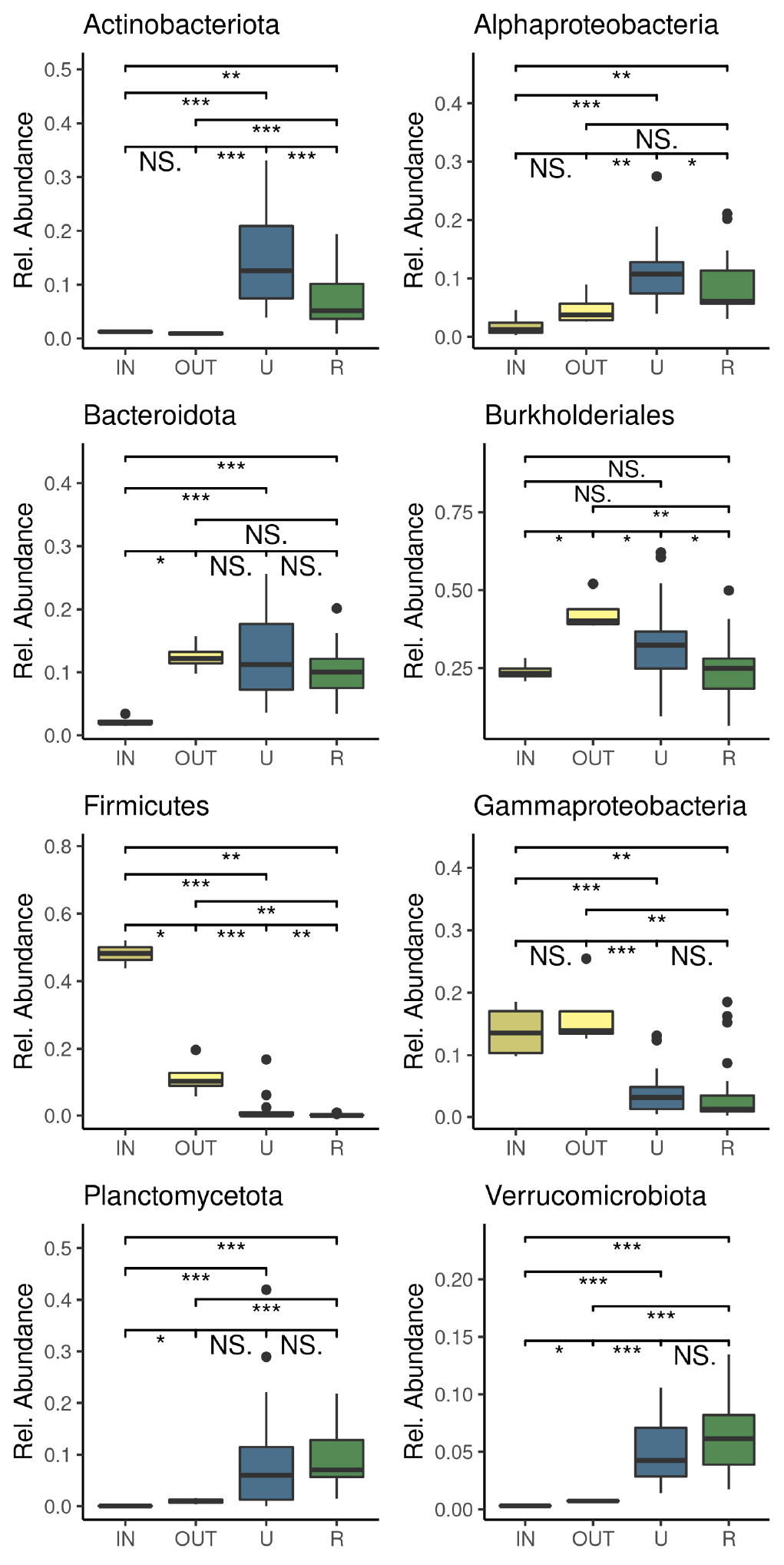
Differences in relative abundance of dominant bacterial phyla between wastewater, urban and rural lakes (summing all seasons and sites). Boxplots show the relative abundance of the most abundant bacterial phyla for wastewater inflow (IN), wastewater outflow (OUT), urban lakes (U: Weisser See, Müggelsee, Feldberger Haussee), and rural lakes (R: Dagowsee, Stechlinsee). Significant differences are indicated by asterisks based on Wilkoxon test. Gammaproteobacteria do not include members of the order Burkholderiales, which have been analysed separately. Rel. abundance – relative abundance, NS – not significant.

Among all defined ASVs from water, sediment and wastewater samples (total = 6712 ASVs) 15.9% were shared between all three habitats, i.e. wastewater, urban and rural lakes (**Fig. 3**). 4.1% of ASVs were unique to wastewater, 5.9% were unique to urban lakes and 7.2% were unique to rural lakes. Wastewater shared 3.9% of the ASVs with urban lakes and only 0.3% with rural lakes, while urban and rural lakes shared 62.7% of ASVs.

**Figure 3:**
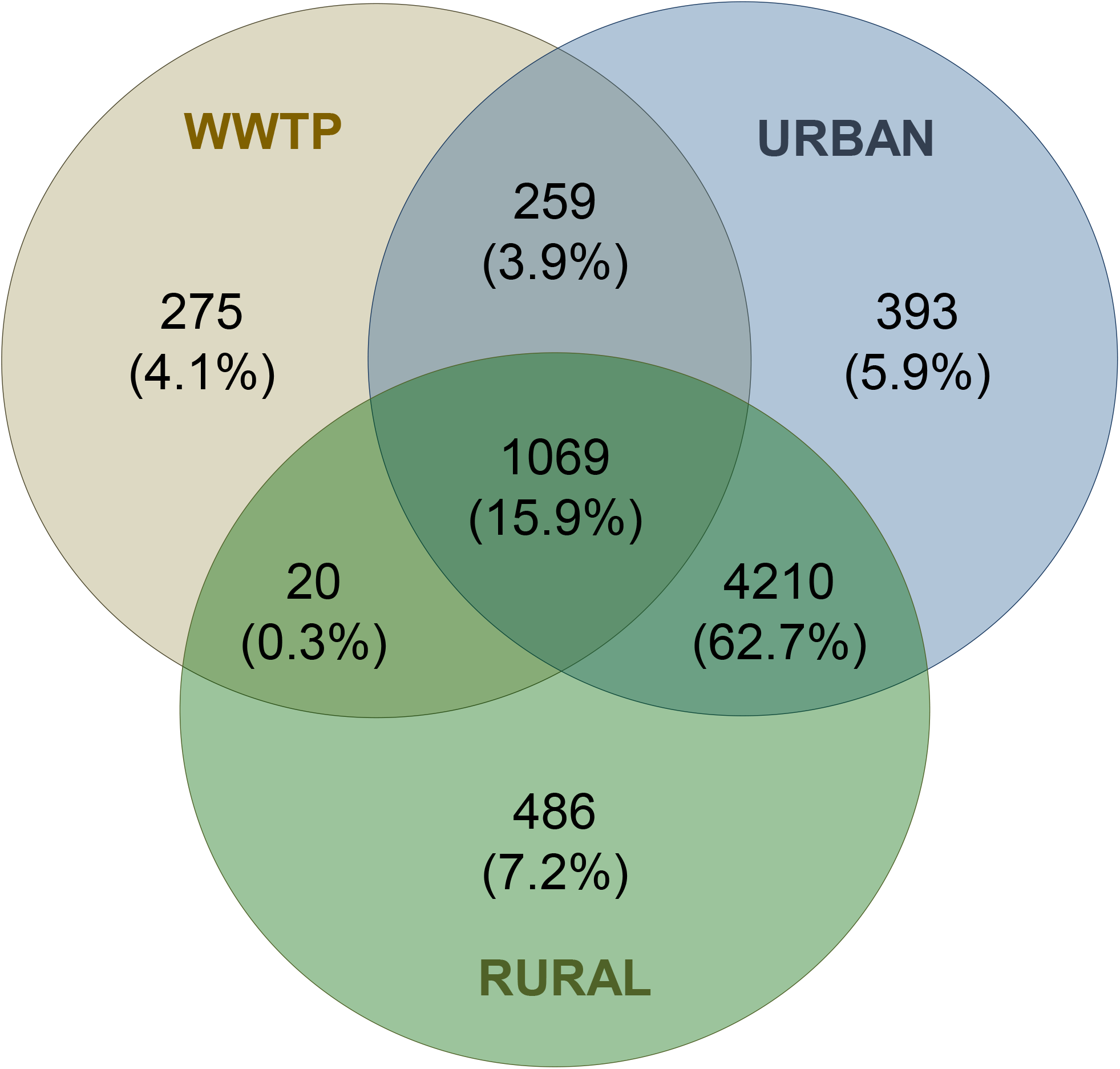
Core, shared and unique ASVs of urban lakes, rural lakes and wastewater. **A** Venn diagram shows core, shared and unique ASVs of the wastewater treatment plant (WWTP inflow and outflow), urban (Weisser See, Müggelsee, Feldberger Haussee) and rural lakes (Dagowsee, Stechlinsee), including sediment samples.

The ternary plots (**Fig. 4)** revealed the distribution of all ASVs of the most abundant bacterial taxa in the three different habitats: rural lake water, urban lake water and wastewater. We excluded the sediment from this analysis to allow for a direct comparison between lake water and wastewater samples.

**Figure 4:**
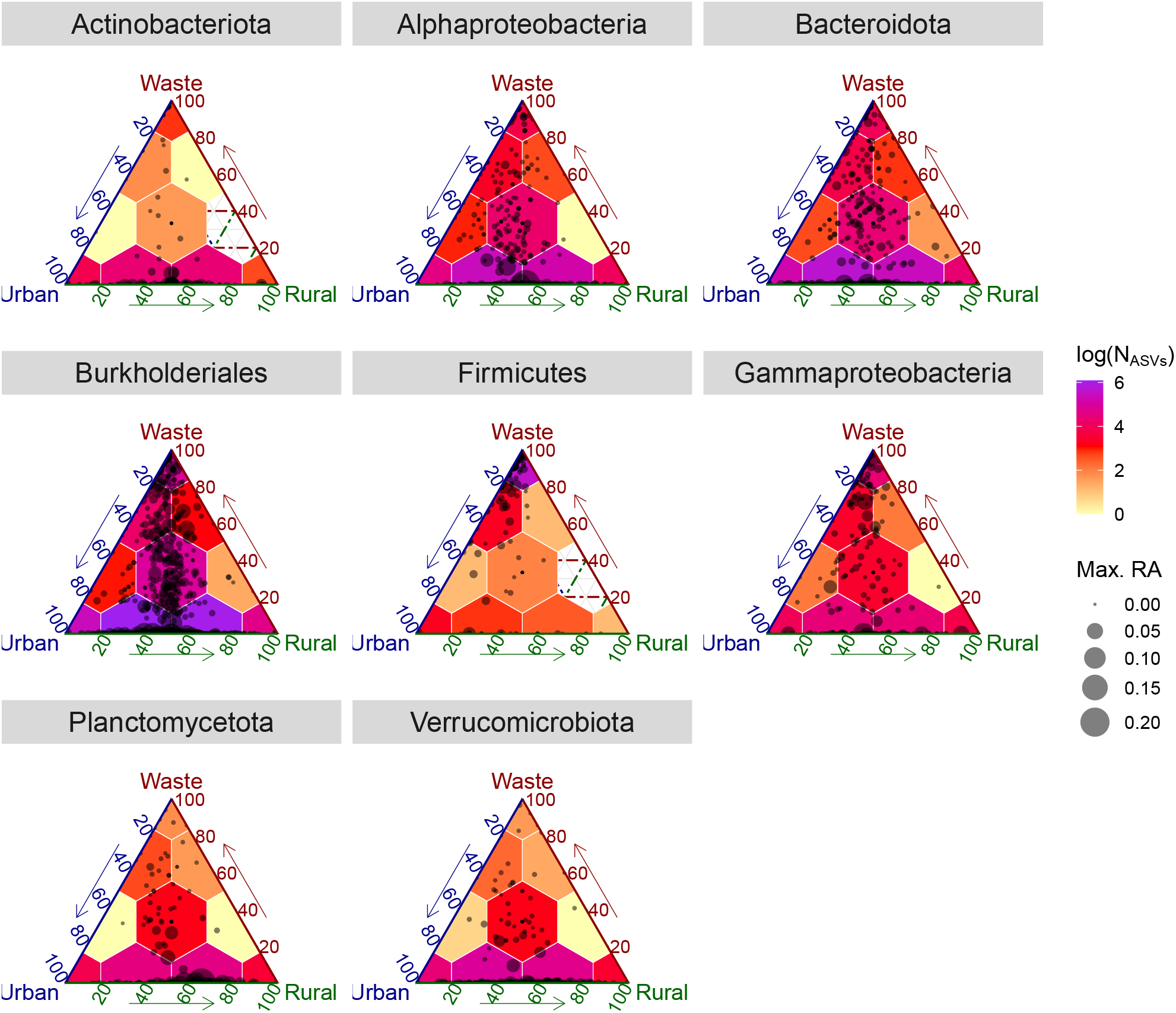
Habitat specific bacterial communities. **[a]** Ternary plots show the number and relative abundance of ASVs (dots) that had 10 or more sequences in at least 3 samples and their occurrence in rural freshwater, urban freshwater and wastewater. Only the most abundant bacterial phyla/groups are shown. Colours in the plots indicate the number of ASVs (log-transformed) and the size of the dots indicate the maximum relative abundance for each ASV. Points close to the corners of the plots represent either ASVs that occur more often or that are specific for that given habitat, while points between two vertexes or in the middle of the plots have similar occurrence or are specific for the combination of the related habitat. Max. RA – maximum relative abundance.

Wastewater and rural lakes shared only very few ASVs among all dominant phyla. Most ASVs were shared between rural and urban lakes, except for the phylum Firmicutes that showed the highest prevalence of unique ASVs in wastewater samples followed by shared ASVs between wastewater and urban lakes (e.g. ASVs belonging to the genera *Acinetobacter, Bacteroides, Bifidobacterium* and *Enterococcus*). The Actinobacteriota and Gammaproteobacteria had several ASVs unique to urban water and several shared between urban and rural lake water. All bacterial phyla showed higher numbers of ASVs in urban lake water than in rural lake water alone. An indicator species analysis (ISA) identified in total 32 ASVs as significant indicators for urban water (urban lakes and wastewater) including the genera *Acidovorax* (Burkholderiales), *Flavobacterium* (Bacteroidota) and *Pseudomonas* (Gammaproteobacteria) (Table C.1).

### 3.3 Bacterial genera that include known potential pathogens

The prevalence of the most relevant genera which include species that are known human pathogens are shown in **Fig. 5a**. While some of the genera such as *Aeromonas, Clostridium* and *Pseudomonas* were equally prevalent in all lakes and wastewater, other genera such as *Alistipes*, *Enterococcus, Escherichia/Shigella* and *Streptococcus* showed a higher prevalence in urban water including wastewater. Furthermore, in some cases the relative abundance was significantly higher in the sediment than in water, for instance for the genera *Bacillus* and *Clostridium* (Fig. A.3). Other taxa such as *Legionella*, *Leptospira*, *Microcystis*, *Mycobacterium* and *Peptoclostridium* had a significantly higher abundance in water samples (Fig. A.3). A weighted correlation network analyses (WGCNA) identified 15 bacterial sub-communities for water and 17 sub-communities for sediment samples that were significantly enriched with bacterial genera containing potential pathogenic species (**Fig. 5b**). Most of these sub-communities did not show a significant correlation with one of the sampled environments, but some of the communities were significantly associated with one specific habitat, e.g. urban waters. The genera *Microcystis, Peptoclostridium* and *Pseudomonas* were enriched in a water sub-community that was significantly associated with urban lakes. The same was true for *Acinetobacter*, *Aeromonas*, *Escherichia*/*Shigella*, *Klebsiella*, *Enterococcus* and *Leptospira*. Sediment sub-communities that were significantly enriched in urban lakes showed a higher prevalence of *Aeromonas, Clostridium, Escherichia/Shigella* and *Streptococcus*. Composition of sub-communities as well as their prevalence in the samples are shown in Fig. A.4. No significant correlation was found between potential pathogenic groups and sub-communities that were only present in rural waters. However, rural sediments harbour sub-communities that favour the presence of *Klebsiella*, *Leptospira*, *Microcystis* and *Pseudomonas*.

**Figure 5:**
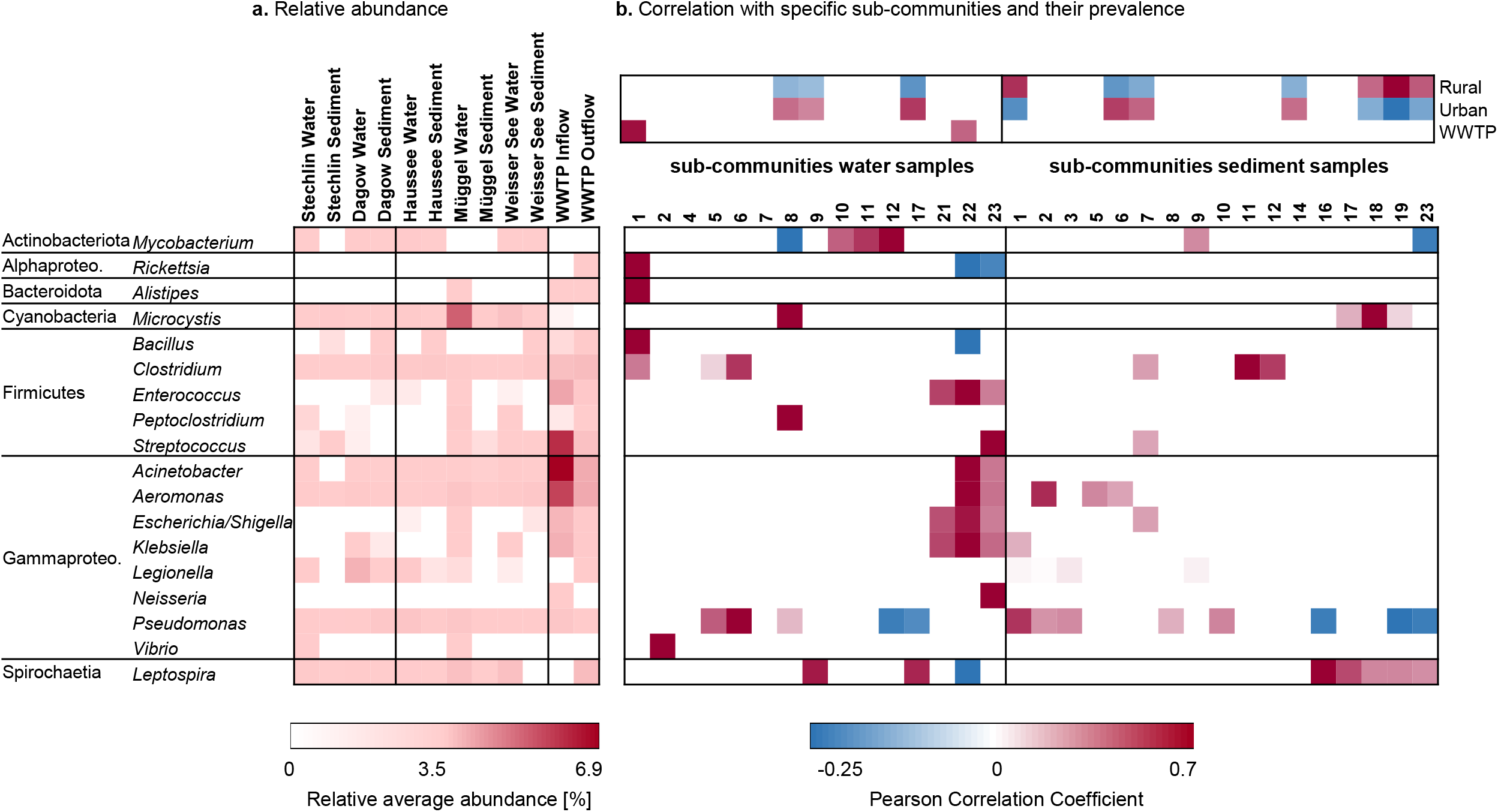
The prevalence of genera that are known to contain potential human pathogens [a] and their correlation with specific sub-communities (1-23) [b]. Heatmap **[a]** shows the average relative abundance in the wastewater treatment plant (WWTP), lake water, and lake sediment. Heatmap **[b]** shows the results of a weighted correlation network analyses (WGCNA). Only significant correlations (p<0.05) of the potential pathogenic genera and specific sub-community structures are shown. The composition and detailed occurrence of the sub-community ASVs can be found in **Suppl. Figure S2**. Alphaproteo - Alphaproteobacteria, Gammaproteo. – Gammaproteobacteria.

A CCA analysis revealed the correlation between potential pathogenic genera (relative abundance) and the measured environmental parameters in lake surface water (**Fig. 6**). All parameters, except nitrite concentration, were significant (one-way ANOVA, p-value 0.002 for nitrate and <0.001 for all other parameters) and, while most bacterial groups were positively correlated with nitrate and temperature, other groups, e.g. *Microcystis* showed a clear correlation with orthophosphate and *Legionella* with DOC.

**Figure 6:**
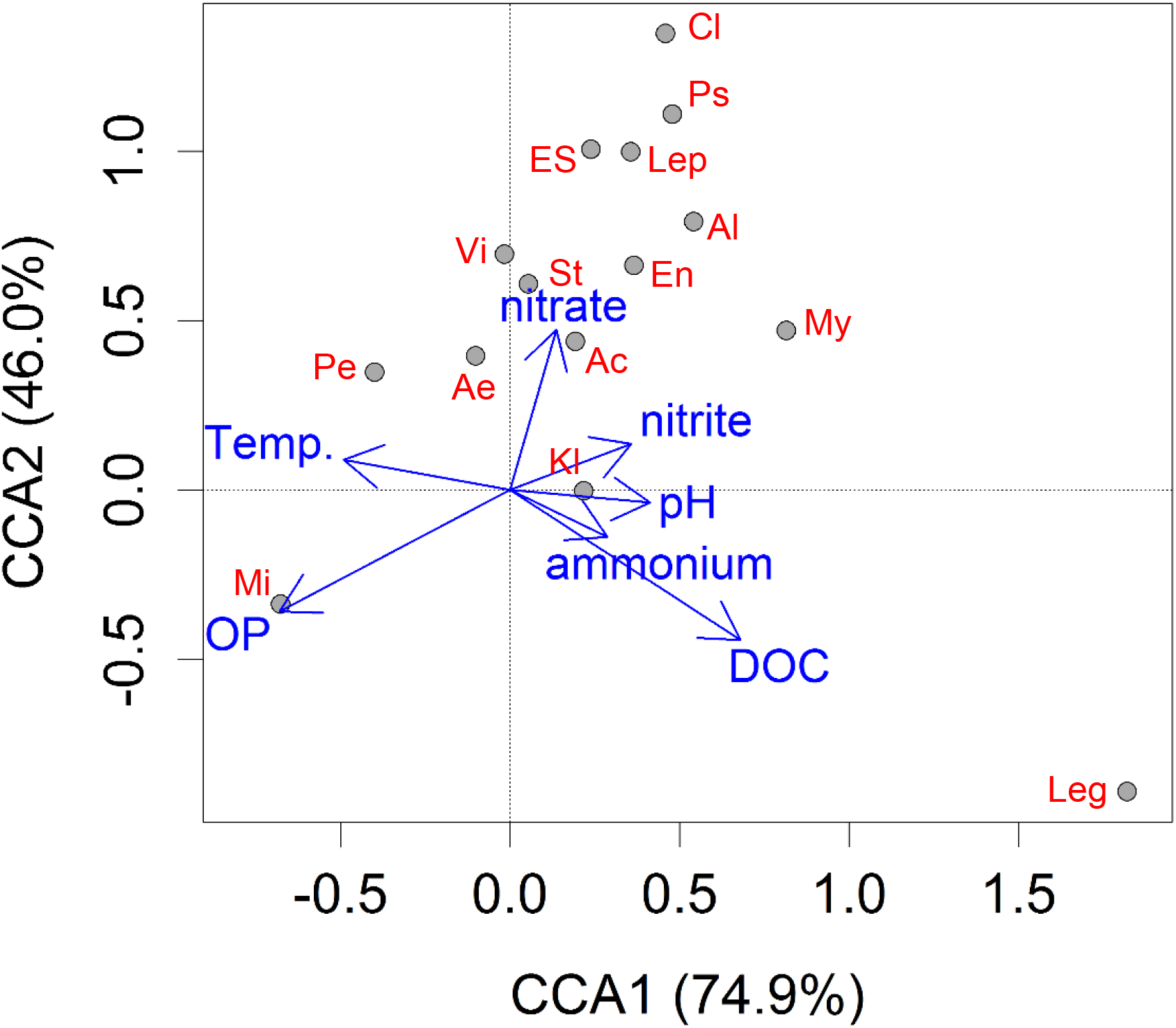
Correlation of potential pathogenic genera and environmental measurements of the water samples. A constrained correspondence analysis (CCA) of genera that are known to contain potential human pathogens from all water samples and the measured environmental measurements: ammonium, dissolved organic carbon (DOC), nitrite, nitrate, orthophosphate (OP), pH, and temperature (Temp.) is shown. *Ac. Acinetobacter, Ae. Aeromonas, Al. Alistipes, Cl. Clostridium (sensu-stricto), En. Enterococcus, ES Escherichia/ Shigella, Kl. Klebsiella, Leg. Legionella, Lep. Leptospira, Mi. Microcystis, Mb. Mycobacterium, Pe. Peptoclostridium, Ps Pseudomonas, St. Streptococcus and Vi. Vibrio.*

## 4. Discussion

Urbanization represents a multifaceted stressor that impacts the quality and function of freshwater systems, promoting eutrophication (Lapointe et al., 2015; Taylor et al., 2004) and contributing to the accumulation of emerging pollutants (Pal et al., 2010; Zinia and Kroeze, 2015). Eutrophication has long been recognised as a major driver affecting microbial community composition. High loads of organic matter lead to increased bacterial activity and creates opportunities for the proliferation of copiotrophs, including many potential pathogens (Smith and Schindler, 2009; Wu, 1999). Eutrophication, however, is not strictly an urban problem. Rural lakes, particularly in close proximity to agricultural land, can also be affected by increased nutrient input. Our results revealed significant differences in the microbial community composition of rural and urban lakes (both water and sediment), and wastewater. Sampled urban lakes, though not directly connected to wastewater effluents, showed a higher similarity to wastewater samples than rural lakes indicating anthropogenic pollution occurs independently of WWTPs. Urban lakes shared 13-fold more ASVs with wastewater than rural lake did (**Fig. 4**). Sewage generally reflects the human faecal microbiome (Newton et al., 2015), suggesting urbanization might have led to a humanization of freshwater bacterial communities (McLellan et al., 2015). In addition, it is known that bathers/swimmers release bacteria from their skin during recreational water activity (Elmir et al., 2007; Plano et al., 2011) and animal or human urine could also be a source of bacterial contamination (Lewis et al., 2013; Rojas et al., 2010).

The presence of habitat specific bacterial communities was supported for wastewater, urban and rural lake water (**Fig. 5a**). The observed differences between rural and urban lake communities appear to be mainly driven by the prevalence of specific dissolved nutrients (**Fig. 2a**). Increased availability of orthophosphate and ammonium coincided with an increase in the relative abundance of most bacterial phyla in urban lakes, particularly the Actinobacteriota, Alphaproteobacteria, Bacteroidota, Firmicutes and Gammaproteobacteria (**Fig. 3**). The diversity (according to the Chao index) was significantly higher in urban lakes for Actinobacteriota, Bacteroidota and Firmicutes (Fig. A4).

Actinobacteriota and Alphaproteobacteria are typically oligotrophic members of freshwater lakes, notably represented by the genera *Planktophila* (acI clade) and *Fonsibacter* (LD12 clade), respectively. These two groups alone can account for up to 50% of the bacterial community composition in lakes in the absence of high phytoplankton biomass (Salcher et al., 2011; Woodhouse et al., 2016). In addition, there is a high capacity for organic matter utilisation within Actinobacteriota and Alphaproteobacteria, in particular by the genera *Planktoluna* (acIV clade) and *Sphingomonas,* respectively (Bagatini et al., 2014). An increased abundance of these latter bacterial taxa in the sampled urban landscapes reflects the enrichment of these copiotrophic taxa at the expense of other oligotrophic taxa. Moreover, the genus *Bifidobacterium* belonging to the phylum Actinobacteriota is a clear indicator of anthropogenic impacts (Bonjoch et al., 2004) and was only found in urban water.

Bacteroidota are well established members of freshwater systems (Newton et al., 2011). They perform important roles in the degradation of organic matter, in particular complex biopolymers (Kirchman, 2002; Newton et al., 2011). Typically, in freshwater systems, dominance and diversity of Bacteroidota are driven by increasing concentrations of either autochthonous (mainly algal or zooplankton biomass) or allochthonous (mainly terrestrial detritus) particulate organic matter. A recent study demonstrates that Bacteroidota strains are highly specific to individual polymeric substrates (Krüger et al., 2019), suggesting that diversity of Bacteroidota scales with diversity of the organic matter pool which may be higher in urban than in rural lakes due to the multitude of different organic matter sources. The greater diversity of Bacteroidota in urban lakes supports this notion (Fig. A5). High terrestrial-aquatic coupling and the dynamic nature of urban landscapes implies a greater diversity of organic matter including faecal contamination (Buccheri et al., 2019; Dick et al., 2005; Krentz et al., 2013) in urban than in rural lakes where cyanobacterial derived autochthonous organic matter seems to be dominant. The genus *Bacteroides,* a known faecal contamination indicator (Hong et al., 2008; Kabiri et al., 2013), was a clear urban signature in our study. Members of Prevotellacae, Rikenellaceae, Tannerellaceae and Weeksellaceae were also significantly enriched in urban waters. In rural lakes the bacterial families Chitinophagaceae, Flavobacteriaceae, Saprospiraceae and Spirosomaceae, well-known freshwater taxa and decomposers of complex carbon sources such as from phytoplankton (McIlroy, 2014; Newton et al., 2011; Raj and Maloy, 1990), were enriched. This suggests there is a multitude and complexity of possible organic matter sources for lakes in urbanized and rural areas.

Firmicutes are not usually abundant in lake water (Newton et al., 2011), but dominate in faeces and wastewater (Buccheri et al., 2019; Newton et al., 2015; Numberger et al., 2019a; Turnbaugh et al., 2007). In our samples, Firmicutes were highly abundant in wastewater, particularly the inflow samples and showed a clear enrichment in urban lakes, but not in rural lakes. Enrichment in urban lakes can be explained by an increase of typical human-derived groups such as Enterococcaceae, Eubacteriaceae, Peptostreptococcaceae, Ruminococcaceae, Streptococcaceae and Veillonellaceae. This “human footprint” also includes genera with potential pathogenic species such as *Acinetobacter, Clostridium, Enterococcus*, *Escherichia*/*Shigella, Klebsiella, Rickettsia* and *Streptococcus*. Previously, toxigenic *C. difficile*, a well-known human pathogen, was isolated from a summer sample of the urban lake “Weisser See” (Numberger et al., 2019b). This supports the hypothesis that urbanization creates and supports favourable bacterial communities promoting the growth of potential pathogens and thus, increases the risk for waterborne or –transmitted bacterial infections.

Gammaproteobacteria generally occur at low abundance in natural freshwater lakes (Lindström and Leskinen, 2002; Newton et al., 2011; Zwart et al., 2002). The increased relative abundance of Gammproteobacteria in urban and rural lakes was due to the abundance of members of Burkholderiaceae, Comamonadaceae and Methylophilaceae, all belonging to the order Burkholderiales. Although the relative abundance of Gammaproteobacteria as a whole did not increase in urban lakes, pronounced urban lake signatures were observed as represented by Aeromonadaceae, Enterobacteriaceae, Moraxellaceae, Pseudomonadaceae, Succinivibrionaceae and Xanthomonadaceae. These bacterial families, enriched in urban waters, include potential human pathogens such as *Aeromonas hydrophila* and *Pseudomonas aeruginosa* constituting a potential health risk. A positive correlation of Gammaproteobacteria with lake eutrophication has been previously observed (Kiersztyn et al., 2019) and relates to the fact that Gammaproteobacteria grow faster than the average lake bacterioplankton, particularly when nitrogen and phosphorus levels are high (Gasol et al., 2002; Šimek et al., 2006). Moreover, our CCA (**Fig. 2b**) showed a clear positive correlation of Gammaproteobacteria with orthophosphate, and to a lesser degree, nitrogen-based nutrients. Although we cannot clearly distinguish between urbanization and eutrophication as a driver for enrichment of Gammaproteobacteria, urbanization leads to an unavoidable eutrophication of aquatic ecosystems (Bowen and Valiela, 2001; Yu et al., 2012) and hence, creates highly favourable conditions for these potential pathogens. In particular, Aeromonadaceae and Pseudomonadaceae have been identified as potential reservoirs for antibiotics resistance genes in various aquatic environments. Thus, they constitute a further potential threat to environmental and human health due to their ability to spread these genes to other harmful microorganisms (Bert et al., 1998; Figueira et al., 2011; Stalder et al., 2019).

Within lakes, bacterial communities were more stable over the four sampled time pointsin sediments than in surface water. Sediment samples revealed a higher bacterial diversity than the water column which has been shown previously (Feng et al., 2009). Sediment provides a more stable environment and can protect microbes, e.g. against environmental changes, UV radiation, drifting and grazing. Furthermore, sediment grains serve as a substrate for microbial biofilms, which further enhance microbial stability and persistence of specific taxa in the system (Haller et al., 2009; Walters et al., 2013). Some bacterial groups that include potential pathogens were also present in urban sediment samples and showed a significantly higher relative abundance than in water, e.g. *Clostridium, Bacillus* and *Pseudomonas* (Fig. A3). This is not surprising since sediments also provide sub-oxic or anoxic conditions which favour many human pathogens such as Enterobacteriaceae (Halda-Alija et al., 2001). Toxigenic *C. difficile* isolated from the sediment of the urban lake ‘Weißer See’ (Numberger et al., 2019b) and other studies demonstrate an extended persistence of faecal indicator bacteria such as *Enterococcus* associated with lake sediments, representing an often neglected reservoir function for human pathogens (Haller et al., 2009; Walters et al., 2013).

In our study, urban lakes contained a higher proportion of taxa which included potentially pathogenic organisms suggesting that urban waters contaminated with pathogenic bacteria enable these bacteria to find a favourable environment in which to proliferate (McLellan et al., 2015; Numberger et al., 2019b; Plano et al., 2011; Walters et al., 2011; Wiedenmann et al., 2006). The presence of bacterial sub-communities, which were mainly present in urban water and significantly enriched with some potential pathogenic groups, highlights this emerging health risk. Nevertheless, while the occurrence of “true” pathogenic species was rare (very low number of sequences) in this study, the enrichment of taxonomic groups to which they belong was constantly present in all urban samples. This increases the risk of stochastic and sudden outbreaks of pathogenic bacteria in urban settings which are less likely in rural settings where less favourable environmental conditions for such copiotrophic, pathogenic bacteria prevail. This potential health risk of urban water bacterial communities may need to be accounted for in future urban lake management strategies (Hipsey and Brookes, 2013; Naselli-Flores, 2008). This also holds true for coastal marine waters in the proximity of wastewater output, where the presence of potential pathogenic groups may increase health risk for waterborne or –transmitted diseases, in particular from urbanized areas (Buccheri et al., 2019).

## 5. Conclusions

Increased urbanization will accelerate “humanization” of aquatic bacterial communities. A better understanding of the ecological and functional consequences of urbanization and the roles of habitat specific bacterial groups is needed to mitigate potential health risks of urban bacterial communities. We identified specific taxa that can exploit ecological niches in urban water (i.e. human-derived bacterial groups such as *Alistipes*, *Bifidobacterium, Bacteroides, Enterococcus, Rickettsia,* and *Streptococcus*), and demonstrate that specific environmental conditions and the presence of specific sub-communities of bacteria promote the emergence and spread of taxa that are known to contain pathogenic species. Urbanization potentially favours aquatic microbiomes supporting the growth of pathogens and antibiotic-resistant bacteria that sporadically enter urban water systems. In contrast, natural waters due to differences in amount and quality of nutrients as well as organic matter and generally lower water temperatures form barriers which greatly limit this health risk. Consequently, urbanization and subsequent humanization should be taken as a serious emerging risk for spreading, propagation and transmission of human pathogens and antibiotic resistance. Beyond the increased proliferation of pathogenic and antibiotic-resistant microorganisms in urban waters, urbanization is likely to have additional impacts on aquatic biodiversity and biogeochemical cycling. Further research is required to explore these little studied and largely unknown impacts of urbanization on aquatic ecosystems. In the frame of the predicted increase in future urbanization, immediate action needs to be taken to mitigate the expected severe impact on aquatic ecosystems including disruptive effects for both humans and environment.

## ACKNOWLEDGEMENTS

We sincerely thank Claudia Quedenau, Michael Sachtleben, Uta Mallock and Elke Mach for technical and Dr. Daniele Y. Sunaga-Franze for computational assistance. This work was supported by a project grant (IRG 3 - Water) from the Leibniz Research Alliance “INFECTIONS’21 - Transmission Control of Infections in the 21st Century” as funded by the Leibniz Association, Germany (SAS-2015-FZB-LFV) and the Leibniz SAW project “MycoLink” (SAW-2014-IGB). The German Federal Ministry of Education and Research (BMBF) supported this project within the Collaborative Project “Bridging in Biodiversity Science - BIBS” (funding number 01LC1501G).

